# Causation without correlation: parasite-mediated frequency-dependent selection and infection prevalence

**DOI:** 10.1101/2021.11.13.468422

**Authors:** Curtis M. Lively, Julie Xu, Frida Ben-Ami

**Author notes:** Author for correspondence. C.M. Lively.

## Abstract

Parasite-mediated selection is thought to maintain host genetic diversity for resistance. We might thus expect to find a strong positive correlation between host genetic diversity and infection prevalence across natural populations. Here we used computer simulations to examine host-parasite coevolution in 20 simi-isolated clonal populations across a broad range of values for both parasite virulence and parasite fecundity. We found that the correlation between host genetic diversity and infection prevalence can be significantly positive for intermediate values of parasite virulence and fecundity. But the correlation can also be weak and statistically non-significant, even when parasite-mediated frequency-dependent selection is the sole force maintaining host diversity. Hence correlational analyses of field populations, while useful, might underestimate the role of parasites in maintaining host diversity.

Subject Area. Evolution

## 1. Introduction

An appealing idea in disease ecology is that coevolving parasites act to maintain genetic diversity in natural host populations [1, 2]. The idea assumes that parasites must mimic host cell-surface molecules in order to evade the host’s immune system [2, 3]. This kind of system favors parasite genotypes that can infect the most common host genotypes, thereby giving an advantage to rare host genotypes [2, 4–8]. Host diversity is thus expected to accumulate due to parasite-mediated, negative frequency-dependent selection. Under this reasoning, it seems sensible to predict that host genetic diversity would be positively correlated with disease prevalence in the wild [e.g., 9, 10].

On the other hand, strong parasite-mediated selection can drive oscillatory dynamics in both host and parasite genotype frequencies [2, 4, 11, 12]. Such dynamics might reduce the association between genetic diversity and parasite prevalence, at least during parts of the coevolutionary cycle. In addition, high host genetic diversity might reduce disease spread, leading to a lower prevalence of infection [13, 14]. This latter idea has support from agricultural studies [15–17], as well as from natural populations [e.g., 18, 19, 20] [recent reviews in, 21, 22].

The potentially complex relationship between (1) negative frequency-dependent selection, (2) genetic diversity, and (3) oscillatory dynamics suggests that the association between genetic diversity and disease prevalence in natural populations might be complex and/or nonintuitive. In the present paper, we used simulation studies to examine when coevolving parasites maintain genetic diversity in clonal host populations, and when one might expect to find a significantly positive correlation between infection prevalence and host genetic diversity. We found that host-parasite coevolution can lead to the maintenance of high levels of host genetic diversity, provided the strength of parasite-mediated selection is stronger than host fecundity selection. We also found regions of parameter space for which the correlation between host diversity and parasite prevalence was positive and statistically significant; but the region is small, and it only partially overlaps with the parameter space for high genetic diversity.

## 2. Simulation Model

We used Excel to simulate host-parasite coevolution in 20 populations. We assumed clonal host reproduction, as neutral genetic markers can be used to infer different multi-locus resistance genotypes from field-collected organisms. This approach is especially helpful when the genetic architecture of resistance is unknown, and it has often been used by field biologists to study natural host-parasite interactions, especially in freshwater snails and waterfleas [e.g., 7, 10, 23, 24]. The hosts were assumed to be haploid, where resistance was determined at two different loci, with three alleles at each locus, giving nine different possible genotypes. Given that the hosts were assumed to be asexual, this set up is the same as that for a single locus with nine alleles. The parasites were also assumed to be haploid asexuals with the same complement of genotypes. We assumed a matching alleles model of infection, meaning that parasites with alleles that did not match their host at both loci were killed by the host’s self-nonself recognition system [25, 26]. Both the hosts and the parasites were assumed to be annuals, meaning that the host and parasites have synchronized life cycles.

The nine host clones were initiated at different randomly selected frequencies in the 20 populations. To simulate fecundity selection among clones, the maximum fecundity for uninfected hosts (*b*_u_) was then randomly assigned to each clone separately in all populations to give a mean within-population fecundity of 10 with a standard deviation of either 2.0 or 1.0. This randomization process meant that the same clonal genotypes would likely have had different maximum fecundities in different populations, which could, for example, be a result of resource differences among populations and/or differences in mutational loads among populations. The process was meant to establish the kind of fecundity differences among clones that is known for freshwater snails and *Daphnia* [e.g., 27, 28]. As expected, fecundity selection in the absence of parasites rapidly eroded clonal host diversity in the simulated populations.

The realized fecundity for uninfected individuals (subscript u) was density-dependent and equal to

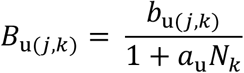

Here *b*_u(*j*,*k*)_ is the number of offspring that would be produced by uninfected individuals of the *j*^th^ clone in the *k*^th^ population in the absence of conspecific competitors; *a*_u_ reflects the sensitivity of uninfected individuals to competition in all populations, and *N_k_* gives the total number of hosts in the *k*^th^ population [following 29]. In the present study, we set *a*_u_ = 0.0001. The carrying capacity (*K*) of an uninfected population, assuming *b*_u_ =10 and *a*_u_ = 0.0001, is *K* = (*b*_u_ − 1)/*a*_u_ = 90,000. The carrying capacity of partially infected populations can be somewhat lower.

The realized fecundity for infected individuals (subscript i) was calculated as a proportion of the realized fecundity of uninfected individuals:

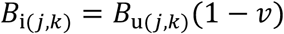

where *v* (for virulence) reflects the reduction in fecundity for infected individuals relative to uninfected individuals. Here *v* = 1 - *b*_i(*j*,*k*)_/*b*_u(*j*,*k*)_, where *b*_i(*j*,*k*)_ is the maximum number of offspring produced by infected individuals of the *j*^th^ clone in the *k*^th^ population. In the simulations, we altered virulence by changing the value for *b*_i_ relative to *b*_u_. Lower values of *b*_i_ reflected higher virulence.

The host populations were all initiated at 90,000 total individuals. To initiate disease spread, an infected adult from each host clone was introduced to all populations at generation 1. In subsequent generations, hosts became infected as juveniles if they contacted one or more genetically matched propagules (spores or eggs) that were shed into the environment by infected hosts in the previous generation. The number of infected juveniles for each host genotype in each population was calculated as,

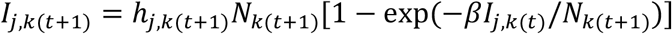

where *β* is the realized parasite fecundity, which gives the number of propagules produced by a single infection that make contact with a host in the next time step. Note that *β* is expected to be a small fraction of the total number of propagules released by infected hosts. *I*_*j,k*(*t*)_ is the number of infected individuals for the *j*^th^ host genotype in population *k* at time *t*; *h*_*j,k*(*t*+1)_ is the frequency of hosts with genotype *j* in population *k* at time *t*+1, and *N*_*k*(*t*+1)_ is the total number of hosts in the *k*^th^ subpopulation. The expression in square brackets gives the probability of infection, assuming that exposure to a single matching parasite genotype is sufficient to cause infection. Infected hosts did not recover from infection.

Similarly, the number of uninfected juveniles for each host genotype *j* in each population *k* was calculated as,

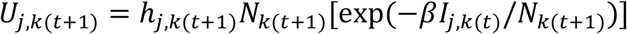

Here the term in square brackets gives the Poisson-distributed probability for the “zero class,” meaning that the host is not exposed to a matching parasite genotype and remains uninfected. This basic approach to modeling infection dynamics was borrowed from previous studies [30, 31]. A recent exploration of the approach is given by [32].

To simulate host and parasite migration, we coded the simulation to introduce a single infected individual from each host clone with a probability of 0.02 in all populations. Hence, a single infected migrant for each clone was introduced, on average, every 50 generations. Similarly, a single uninfected migrant for each host clone was introduce with a probability of 0.05 (giving an average of one migrant every 20 generations). The migrants were assumed to be drawn from a very large metapopulation, of which the 20 simulated populations were only a small part. The main purpose was to introduce genetic variation into the parasite population. In the absence of migration, parasite genetic diversity could be stochastically lost. This is especially true for highly virulent parasites, which tend to generate high-amplitude oscillations that push allele frequencies near to fixation during parts of the cycle [33] (Fig. 2). A total loss of parasite alleles would further decouple the correlation between infection prevalence and host genetic diversity. As such, the incorporation of a low level of migration to restore diversity is conservative with respect to the findings reported here.

The simulation was run for 2100 generations, and the data from each of the 20 populations were collected at the final generation. This was meant to capture the system at a single time point in the way in which field biologists might sample a metapopulations. At generation 2100, we calculated the prevalence of infection for each population. We also calculated host genotypic diversity within each population as

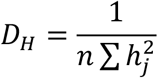

where *n* gives the number of possible genotypes in the population (here *n* = 9), and *hj* gives the frequency of the *j*^th^ host clone. This measure gives the standardized inverse Simpson’s index, with a maximum value of 1.0 and a minimum value of 1/*n* [34]. We also calculated the mean prevalence and mean *D*_H_ across the 20 subpopulations, as well as the correlation between infection prevalence and *D*_H_ across the subpopulations. We iterated the simulation for a range of values for parasite fecundity (*β* = 2, 4, 6, 8, 9, 10, 12, 15, 18, 21, 24, 27, 30, 33, 36) across a range of virulence values (*v*) from 0.10 to 0.90 in increments of 0.10. Using a standard deviation for average host fecundity equal to 2.0, the 2100-generation simulation was run 1000 different times for all combinations of values. The data presented are the averages of these 1000 runs for the final generation (generation 2100) (Fig. 1). We repeated the whole process after reducing the standard deviation for host fecundity from 2.0 to 1.0, which reduced the effect of fecundity selection. To reduce the processing time, we also reduced the number of iterations of the 2100-generation simulation from 1000 to 100.

**Figure 1.**
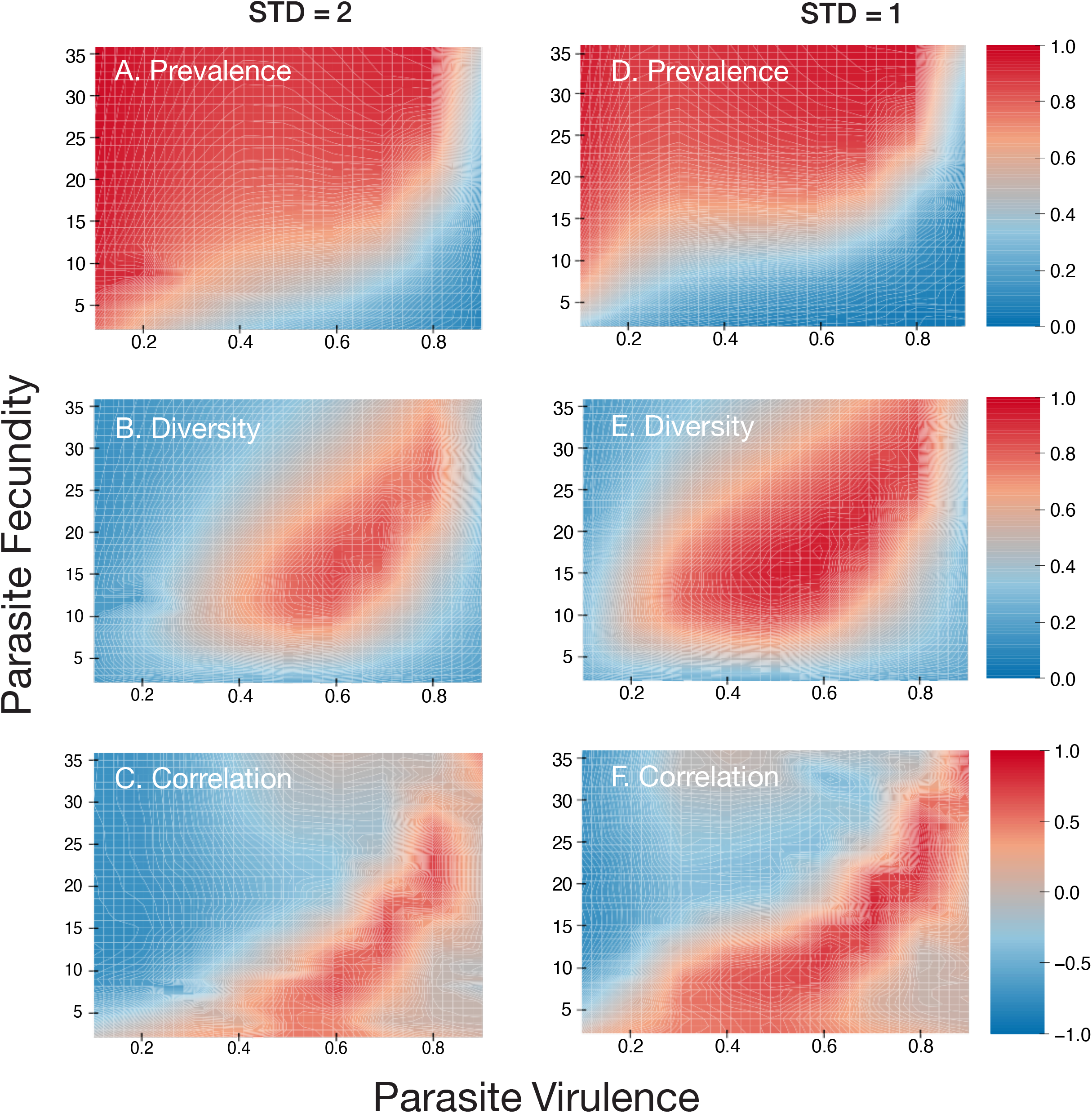
Simulation results. The left-hand column (A-C) gives simulation results for which the standard deviation in host fecundity was 2.0, while the right-hand column (D-F) gives results for a standard deviation in host fecundity of 1.0. The higher standard deviation (A-C) gives stronger fecundity selection. The top row (A & D) gives the results for mean infection. The middle row (B & E) gives the results for mean diversity, while the lowest row (C & E) gives the correlation between diversity and prevalence of infection. The side bars show the values of the different colors for prevalence (A & D), diversity (B & E), and the correlation between diversity and prevalence. For graphs C and E, correlations having an absolute value of greater than 0.45 have *P* values less than 0.05. Note that, as expected, high levels of diversity are maintained across more of the parameter space when fecundity selection is weaker. Note also that high correlations are shifted to the South and West relative to regions of high host diversity. As such, high levels of host diversity can be associated with low (or even negative) correlations between diversity and prevalence. Note that the simulation results assumed clonal reproduction, but they could also be applicable to highly inbred populations.

**Figure 2.**
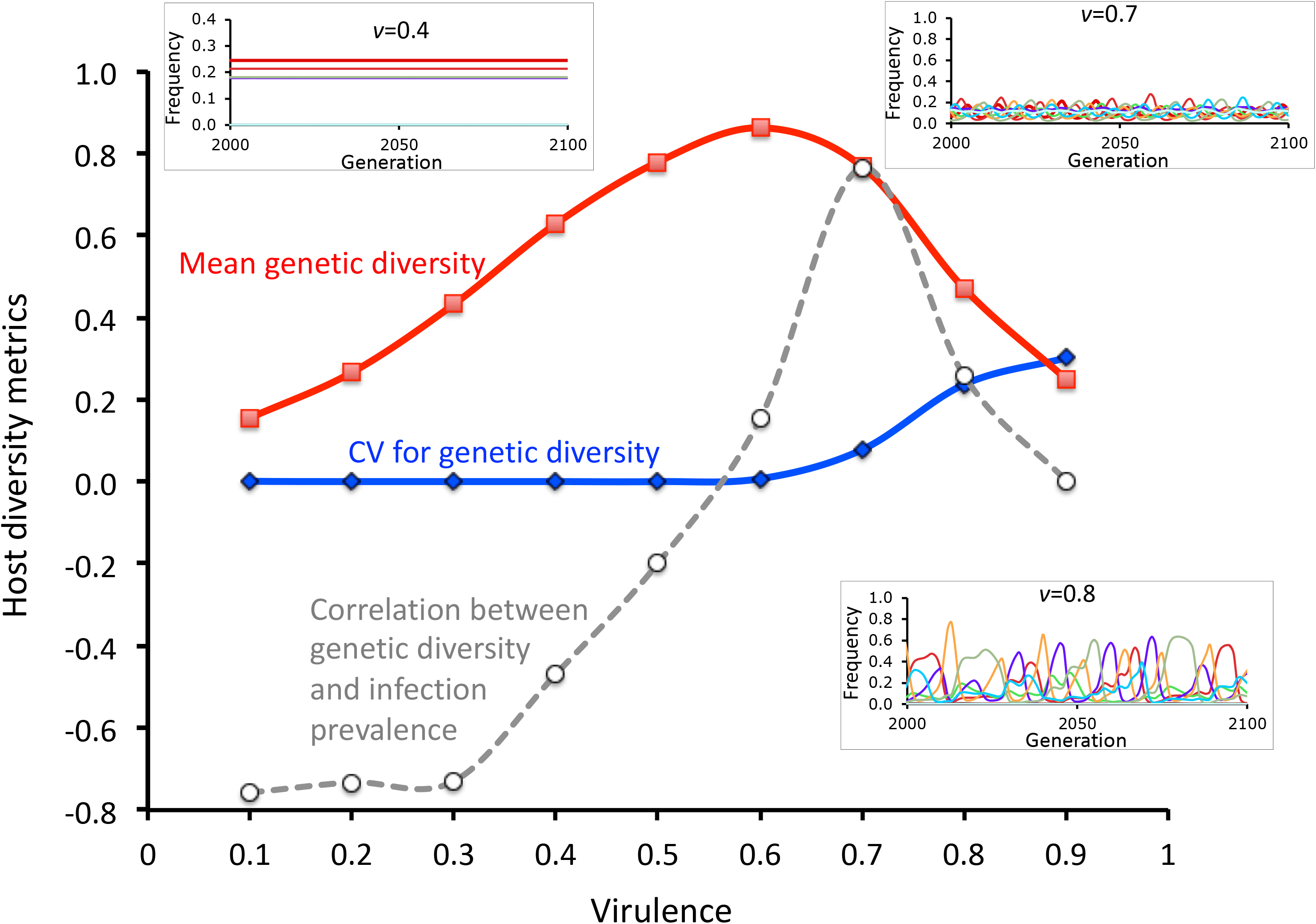
Cross section through Figs 1 B & C at parasite fecundity = 15. The lines show the relationships between host genetic diversity (red line), fluctuations in diversity (blue line, the coefficient of variation for host diversity), and the correlation between host genetic diversity and infection prevalence (dashed grey line). Note that the correlation is only positive and high for a small portion of parameter space in which diversity is high. Also note that the peak correlation corresponds to moderate fluctuations in diversity over time (grey line). Insert, upper left: representative host genotype dynamics for moderate virulence (*v* = 0.4). The different colored lines represent different host genotypes. Insert, upper right: representative host genotype dynamics for moderately high virulence (*v* = 0.7). Insert, lower right: representative host genotype dynamics for high virulence (*v* = 0.8). Taken together, the results suggest that detection of strong positive correlations would be most likely at moderate-to-high levels of parasite-mediated selection, depending on parasite fecundity (Fig. 1).

## 3. Results and Discussion

Our main goal was to determine the effects of parasite fecundity and parasite virulence on host genetic diversity and infection prevalence. Not surprisingly, the mean prevalence of infection increased with parasite fecundity, especially when parasite virulence was low (Fig. 1A,D). However, for higher levels of parasite virulence, the prevalence of infection decreased (Fig. 1A,D). This decrease in prevalence with higher levels of virulence does not stem from the damping of infection, such as that observed in virulent contagious diseases [e.g., 35, 36]. In our simulation models, the infection is not contagiously transferred among hosts, but is rather randomly distributed among juvenile hosts in the next generation. The reduction in disease is because higher levels of host genetic diversity were maintained with higher levels of parasite virulence (virulence = 0.4-0.8). This result can be seen by comparing Figs. 1A and 1B: the transition from high to low prevalence (red to blue) in Fig. 1A occurs where genetic diversity is increasing (blue to red in Fig 1B).

Why does host-genetic diversity increase for virulence levels between roughly 0.4 and 0.8? The simplest explanation is that parasite-mediated selection becomes strong enough in this region to counter the effects of fecundity selection. Given that the nine host clones were initiated with variable intrinsic birth rates, the most fecund clone eliminates all other clones in the absence of parasites (clonal selection). Hence, for parasites having low virulence, host genetic diversity is eroded by clonal selection, leading to higher levels of infection in the genetically depauperate host population. As virulence increases, parasite-mediated frequency-dependent selection leads to higher levels of host genetic diversity, because selection against the more common and more fecund host clones counters any fecundity advantage that they have. Hence, more virulent diseases lead to higher levels of host genetic diversity, which reduces the overall prevalence of infection. Also note that the most common clones are most fecund when uninfected, but also the most infected. This pattern would look like a trade-off in which fecundity trades off with resistance. But there is no inherent trade-off. The most common local clones would not be more infected by allopatric parasites, which are adapted to a different set of host genotypes.

Curiously, however, both genetic diversity and prevalence of infection decline as the parasites become very virulent (*v* > 0.8). This leads to the hump-shaped topography observed for the effects of virulence and parasite fecundity on host diversity (Fig. 1B,E). To understand the pattern, we ran a “transect” through the hump. Specifically, we held realized parasite fecundity constant at 15, which meant that each infection produces 15 propagules that contact a host (even though 15 is likely to be a very small fraction of the total propagules produced). We then evaluated the effects of different levels of virulence on genotype frequency dynamics using the same simulation model as described above. To do this, we calculated the mean CV (coefficient of variance) for clonal frequencies for the last 100 generations of the simulation. We then took the average of these values over all 20 populations. High average values would be indicative of high amplitude oscillations in genotype frequencies over time.

The “transect” through the surface reflects the hump shaped pattern for genetic diversity (plotted against virulence) shown in Fig. 2. It also shows that the genotypic oscillations increased with virulence for values of virulence greater than 0.6. In addition, mean diversity decreased with increasing amplitude in genotype-frequency oscillations (Fig. 2), which was a direct result of different genotypes becoming periodically very common over time. Thus, even though negative frequency-dependent selection is maintaining variation in each of the host populations over time, the diversity in the host population can periodically be relatively low.

Finally, we examined the correlation between prevalence of infection and genetic diversity (Fig 1 C,F). Our goal here was to determine the likelihood of detecting patterns in nature when, in fact, parasites are a primary factor in maintaining host genetic diversity. The results suggest that positive and significant correlations can be observed in some parts of the parameter space, but this region of parameter space only partially overlaps with the region of highest host genetic diversity (compare Figs 1B and 1C). Hence, our results suggest that the lack of significance, or even slightly negative correlations between infection prevalence and genetic diversity, cannot be taken as evidence that parasites are unimportant to the maintenance of diversity in host populations. Perhaps analogously, the lack of significant correlations between the frequency of sexual females and the prevalence of infection in mixed (sexual and asexual) populations of hosts may not be a reliable signal that parasites are unimportant in selection for cross-fertilization.

## 4. Summary

We conducted a simulation study of host-parasite coevolution in a metapopulation of clonal hosts to better understand the relationship between host genetic diversity and prevalence of infection. In our model, the only force maintaining genetic diversity was parasite-mediated frequency-dependent selection against common host genotypes. Nonetheless, even in regions of parameter space showing high genetic diversity, the correlation between prevalence of infection and genetic diversity could be so weak as to be unlikely to be detected as statistically significant. Hence, over much of the parameter space, causation does not produce significant correlation. These results suggest that field studies could lead to the false conclusion that parasites are unimportant in selecting for rare genotypes, when in fact parasite-mediated selection is the only selective force leading to diversity.

## Data accessibility

The simulation and representative data are available from Dryad [37]

## Authors’ Contributions

CML and FB-A conceived the project. CML wrote the basic simulation, and FB-A wrote the excel macro. JX ran the simulations, organized the data, and constructed the figures. CML and JX wrote the first draft of the paper, and FB-A and CML revised the paper.

## Acknowledgements

We thank Mandy Gibson, Zoe Dinges, and three anonymous reviewers whose constructive suggestions helped us to improve the presentation.

